# Manipulating the microbiome alters regenerative outcomes in *Xenopus laevis* tadpoles via lipopolysaccharide signalling

**DOI:** 10.1101/2021.12.09.472019

**Authors:** Phoebe A. Chapman, Campbell B. Gilbert, Thomas J. Devine, Daniel T. Hudson, Joanna Ward, Xochitl C. Morgan, Caroline W. Beck

## Abstract

*Xenopus laevis* tadpoles can regenerate functional tails, containing spinal cord, notochord, muscle, fin, blood vessels and nerves, except for a brief refractory period at around one week of age. At this stage, amputation of the tadpole’s tail may either result in scarless wound healing, or the activation of a regeneration programme, which replaces the lost tissues. We recently demonstrated a link between bacterial lipopolysaccharides and successful tail regeneration in refractory stage tadpoles, and proposed that this could result from lipopolysaccharides binding to Toll-like receptor 4 (TLR4). Here, we have used 16S rRNA sequencing to show that the tadpole skin microbiome is highly variable between sibships and that the community can be altered by raising embryos in the antibiotic gentamicin. Six gram-negative genera, including *Delftia and Chryseobacterium*, were over-represented in tadpoles that underwent tail regeneration. Lipopolysaccharides purified from a commensal *Chryseobacterium spp. XDS4*, an exogenous *Delftia spp*. or *Escherichia coli* could significantly increase the number of antibiotic-raised tadpoles that attempted regeneration. Conversely, the quality of regeneration was impaired in native-raised tadpoles exposed to the antagonistic lipopolysaccharide of *Rhodobacter sphaeroides*. Editing TLR4 using CRISPR/Cas9 also reduced regeneration quality, but not quantity, at the level of the cohort. However, we found that the editing level of individual tadpoles was a poor predictor of regenerative outcome. In conclusion, our results suggest that variable regeneration in refractory stage tadpoles depends at least in part on the skin microbiome and lipopolysaccharide signalling, but that signalling via TLR4 cannot account for all of this effect.

## 1. INTRODUCTION

Tadpole tail regeneration in *Xenopus laevis* provides a useful model to study regenerative mechanisms in complex tissues. Tails contain midline neural tube - the forerunner of the spinal cord - as well as notochord, paraxial muscles (somites), blood vessels, nerves and the dorsal and ventral fins (extensions of the epidermis). *Xenopus laevis* is a well-used model organism, and development has been classified into 66 stages, with pre-feeding stages that are well synchronised (Nieuwkoop and Faber 1967). Tails regenerate well following partial amputation from stage 40-44. From stage 45-47, there is a dramatic reduction in the number of tadpoles undergoing regeneration, with a scarless wound healing programme replacing this in many tadpoles (Beck et al. 2003). We refer to this as the refractory period, and it is useful since it offers the opportunity for both gain and loss of function experiments in a single system. Prior studies have implicated many developmental signalling pathways, as well as processes such as apoptosis, epigenetic regulation, membrane depolarisation, extracellular matrix remodelling, reactive oxygen species production, inflammatory response, and metabolic reprogramming in *Xenopus* tail regeneration (for recent review, see (Phipps et al. 2020).

Tails in the refractory period seem to commit to either regeneration or wound healing pathways in the first six hours following amputation (Beck et al. 2003). Tails that successfully recruit regeneration organising cells (ROCs) to the wound site to form a wound epithelium will go on to organise the regeneration of either fully patterned or pattern-deficient tails (Aztekin et al. 2019) via recruitment of underlying distal cells to a regeneration bud (Slack et al. 2004). Tails that instead heal with a full-thickness epidermis, including a basement membrane, will not regenerate, and do not form a regeneration bud (Beck et al. 2009). In many regeneration competent model organisms, macrophages (phagocytic cells that form part of the innate immune system) are critical for regeneration. This is true of zebrafish tails (Li et al. 2012; Petrie et al. 2014) and axolotl limbs and heart (Godwin et al. 2013; Godwin et al. 2017), as well as *Xenopus* tadpole tails (Aztekin et al. 2020). Recent work from our lab has shown that the base rate of tadpole tail regeneration is innately variable, with some sibships showing naturally higher regenerative rates during the refractory period (Bishop and Beck 2021). Raising tadpoles the presence of aminoglycoside antibiotics, which is often done prophylactically in labs and would be expected to alter the microbiome, reduces the percentage of regenerators in a cohort (Bishop and Beck 2021), suggesting that the microbiome may be important in refractory period regeneration efficiency. Microbiomes are important for wound healing in a lot of model animal systems, including planarians (Arnold et al. 2016; Williams et al. 2020) and mice (Velasco et al. 2021; Wang et al. 2021). Among these examples, indole (an aromatic amino acid metabolite produced by gut bacteria) (Williams et al. 2020) and the inflammatory cytokine IL-1β (Wang et al. 2021) have been implicated as critical components of the signalling pathway leading to regeneration.

In *Xenopus*, regeneration in antibiotic treated tadpoles can be returned to baseline levels by exposing the cut tail surface to heat-killed gram-negative bacteria or purified lipopolysaccharides (LPS) (Bishop and Beck 2021). We hypothesised that TLR4, a Toll-like receptor of the innate immune system that recognises LPS (Chow et al. 1999), is exposed to skin bacterial LPS of tadpoles only when the tail is cut (Bishop and Beck 2021). LPS binding of TLR4 on either tissue resident mesenchymal stem cells (Munir et al. 2020) or macrophages (Chow et al. 1999) could produce an inflammatory cytokine response, generating a pro-regenerative environment.

Under laboratory conditions, the most likely source of LPS that could influence tail regeneration is from commensal gram-negative bacteria on the tadpole skin. Here, we have used 16S ribosomal RNA amplicon sequencing to compare tail skin microbiome composition and regeneration success in three sibships (sibling cohorts) of tadpoles, raised with and without antibiotic gentamicin to disrupt gram-negative bacterial flora. We also tested the hypothesis that LPS binding to TLR4 elicits a regeneration response, using both an antagonistic LPS purified from *Rhodobacter sphaeroides* (recently renamed as *Cereibacter sphaeroides* (Hordt et al. 2020)), and gene editing of *Tlr4*.*S*.

## 2. METHODS

### 2.1 Animal ethics

Procedures for production of *X. laevis* eggs and embryos were approved by the University of Otago’s Animal Ethics Committee as AUP19-01.

### 2.2 Animal husbandry

Adult *X. laevis* used in this study are housed within a recirculating aquarium system within PC2 facilities at the University of Otago. The system is supplied with carbon-filtered mains water and frogs are fed twice weekly with salmon pellets. The colony was established in 2004 and has been closed, with no contact with outside animals, since then. Current adults are F_1_ or F_2_ captive bred.

### 2.3 Egg collection and fertilisation

All eggs and embryos used in this work were produced by inducing egg laying in adult female *X. laevis*, weighing 50 to 100 g, by injecting 500 U of HCG (Chorulon) per 75 g of bodyweight into the dorsal lymph sac. Adult males were killed by immersion in a lethal dose of benzocaine. Eggs were laid into 1 x MMR (Marc’s modified ringers, pH 7.4: 100 mM NaCl, 2 mM KCl, 1 mM MgSO_4_.7H_2_O, 2 mM CaCl_2_, 5 mM HEPES, 0.1 mM EDTA at pH 8.0) and fertilised in vitro using 50 μl of fresh male *X. laevis* testes, prepared by lightly disrupting tissue using a plastic pestle to release sperm into 1 ml of MMR. Embryos and tadpoles were raised at 18 °C in an incubator.

### 2.4 Tail regeneration assays

For both the antibiotic treatment and CRISPR/Cas9 editing experiments, groups of tadpoles were raised in 10 mm petri dishes containing 30 ml 0.1 x MMR. For the treatment experiment, tadpoles were raised with or without 50 μg/ml added gentamicin according to treatment group. CRISPR/Cas9 edited tadpoles were raised without gentamicin. Gentamicin was kept constant by adding fresh medium to applicable dishes every second day, and was discontinued 1 day post amputation, by which time wound healing is complete. Tail regeneration assays were done at stage 46 (Nieuwkoop and Faber 1967), in the refractory period (Beck et al. 2003) before commencement of feeding. Tadpoles were immobilised using 1/4000 w/v MS222 (tricaine, Sigma) in 0.1 x MMR and the distal third of the tail was removed using a sterile scalpel blade. Tadpoles were rinsed in 0.1 x MMR to remove MS222. For treatment experiments, tadpole groups were placed back into petri dishes containing their respective media. For gene editing experiments, individual tadpoles were placed into 24-well culture plates in 1 ml 0.1 x MMR, and tail tips were kept for genotyping. Tadpoles were not fed. Tails were scored for regeneration after 7 days as one of four categories: FR (full regeneration, no visible defect, scores 10/10); PG (partial good, tail regenerated but may have a missing fin on one side, or a bend in the tail scores 6.6/10); PB (partial bad, at least one core tissue missing, short, often bent or grows along the ventral fin cut site, scores 3.3); or NR (no regeneration, full-thickness epidermis forms over wound site, scores 0). This is based on the method devised by Adams et al., (2007)(Adams et al. 2007). Scoring using this criteria was done on an unblinded basis by a single person across each experiment, and examples are shown in Figure S1.

To assess the ability of LPS to “rescue” regeneration in gentamicin raised tadpoles, 50 μg/ml or higher of 200 x LPS stock was added to tadpole media after tail amputation and rinsing. Tadpoles were incubated in the LPS solution for 1 hour before being returned to fresh 0.1 x MMR. TLR4 antagonist LPS from *R. sphaeroides* was added for 1 hour post amputation in tadpoles raised with no antibiotics.

### 2.5 Microbial sampling

Tadpole tail samples (stage 46) were acquired by collecting freshly cut tail tips (posterior third of the anatomical tail) from regeneration assays into 0.2 μl 8-strip PCR tubes, adding 50 μl of filter-sterilised sodium chloride / Tween solution (0.15 M NaCl, 0.1% Tween20), and vortexing for 1 minute before storing at −20 °C. Negative controls were generated using the same technique but without adding a tail tip. Ninety-six tadpoles (48 gentamicin-raised and 48 untreated) were collected from each of three sibships. The tadpoles were arrayed in 24-well plates with 1 ml 0.1 x MMR, incubated at 22 °C and assayed for regeneration after 7 days. Tail regeneration was scored as described above, except that the PG and PB regenerates were both classified as “Partial”.

### 2.6 Tadpole microbial culture assay

A qualitative assay was devised to demonstrate the effect of raising tadpoles in gentamicin on the number of viable bacteria on stage 47 tadpole skin. Individual tadpoles from a single sibship (raised with or without gentamicin) were first washed twice in sterile 0.1 x MMR and then vortexed for 20 seconds in 100 μl of sodium chloride / Tween solution. Fifty microlitres of the resulting solution was added to 1 ml of Luria Broth (LB), diluted 10 fold in LB, and spread onto replicate LB agar plates. Plates were incubated at 18 °C for 66 hours and photographed on a black background.

### 2.7 Bacterial culturing

*Escherichia coli* DH10B strain were grown from glycerol stocks at 37 °C in LB overnight with shaking. Commensal bacteria (*Chryseobacterium spp*.) were cultured from adult female *X. laevis* using gentle swabbing of dorsal, ventral and limb skin for a total of 15 seconds with sterile cotton-tipped swabs (Puritan). Swabs were plated onto Oxoid nutrient agar and incubated at 30 °C for 48 hours. Colonies were purified by streaking. Two additional bacterial strains were obtained from culture collections in order to characterise the effects of their LPS: *Delftia Wen et al 1999* (ICMP 19763) was obtained from Manaaki Whenua - Landcare Research NZ (Wen et al. 1999) and *Rhodobacter sphaeroides* (DSM-158, recently reclassified as *Cereibacter sphaeroides*(Hordt et al. 2020)) was obtained from DSMZ (German Collection of Microorganisms and Cell Cultures). Both were grown on Oxoid nutrient agar and incubated at 30 °C.

The identity of the commensal *Chryseobacterium spp*. isolate was determined by whole genome sequencing and ANI analysis, using the same methods described by Hudson et al. (Hudson et al. 2021). The isolate was most closely related to *Chryseobacterium sp. MYb7* (ANI 96.7 %), and has been deposited in the Manaaki Whenua Landcare Research culture collection as *Chryseobacterium* XDS4 (ICMP 24359). It is hereafter referred to as *Chryseobacterium spp. XDS4*.

### 2.8 LPS extraction from gram-negative cultures

Purified bacterial isolates were cultured in Oxoid nutrient broth, grown overnight at 30 °C, heat-killed at 60 °C for 60 minutes, and pelleted by centrifugation at 5000 g for 10 minutes. Pellets were resuspended in 10 ml PBS pH 7.2, re-spun and re-suspended, and pelleted a final time. Pellets were then frozen at −80 °C for at least two hours before freeze drying in a VaO2 vacuum chamber at −80 °C overnight. LPS was extracted from heat-killed and lyophilised bacteria as described by Yi and Hackett (2000)(Yi and Hackett 2000), using TRI-reagent (Sigma). Briefly, each batch used 10 mg lyophilised bacteria and 200 μl TRI reagent in 1.5 ml Eppendorf tubes. LPS was extracted into the aqueous phase with chloroform, and the organic phase was washed 3 x to maximise yield. Nucleotides were removed by 10 U DNAse and 20 μg RNaseA treatment for 10 mins at 37 °C, followed by 20 μg Proteinase K to remove protein and inactivate nucleases for a further 10 mins. Samples were dried in an Eppendorf concentrator plus Speedvac overnight. Finally, LPS pellets were resuspended in cold 500 μl 0.375 M MgCl in 95% ethanol according to Darveau and Hancock (Darveau and Hancock 1983), precipitated at −30 °C for 30 minutes, repelleted at 12000 g for 15 minutes at 4 °C, dried briefly, resuspended in 200 μl ultrapure water, and stored as aliquots at −30 °C. The estimated concentration of 10 mg/ml was based on a 20% yield of LPS from lyophilised bacteria (Yi and Hackett 2000). LPS was checked by acrylamide gel electrophoresis using a BioRad mini Protean and silver staining (Pierce) according to Laemmli (Laemmli 1970). Duplicate gels were stained with 0.5 % Coomassie brilliant blue R250 (Sigma) to confirm no protein. Size was approximated using 5 μl Novex sharp protein marker.

### 2.9 DNA extraction, 16S rRNA amplicon sequencing, and analysis of tail samples

DNA from tadpole tails was extracted using a DNeasy PowerLyser PowerSoil DNA extraction kit (Qiagen) according to the manufacturer’s instructions, eluted into a final volume of 30 μl and stored at −80 °C. Amplification and sequencing of the V4 hypervariable region of 16S rRNA gene by Illumina MiSeq were performed for as described previously by Caporaso et al. (2011) using primers 515F/862R (Caporaso et al. 2011). Sequencing of 229 samples was done at Argonne National Laboratory, Illinois, USA, and used peptide nucleic acid (PNA) PCR clamps to inhibit the amplification of host mitochondrial sequences (Lundberg et al. 2013). Amplicon sequences (2 × 250 bp) were processed using the DADA2 package (version 1.6.0) in R(Callahan et al. 2016) according to authors’ recommended best practices. The taxonomy was annotated using the naïve Bayesian classifier method with the Silva reference database version 128 (Quast et al. 2013). Downstream analyses were performed using R (version 3.4.3), packages vegan (version 2.4.6) (Oksanen et al. 2019) and phyloseq (version 1.22.3) (McMurdie and Holmes 2013). Samples with fewer than 1500 reads were excluded from further analysis. Sequence data for all samples has been deposited with NCBI (BioProject ID PRJNA780297)

### 2.10 CRISPR/Cas9 targeting of *Tlr4*.*S*

ChopChop v2 (Labun et al. 2016) was used to identify four unique sgRNA sequences from *X. laevis Tlr4*.*S* (Table 1). EnGen sgRNA oligo designer v1.12.1 tool (NEB) was used to generate 55 bp oligos. These were synthesised by IDT and converted into sgRNAs using the EnGen Cas9 sgRNA kit (NEB) according to instructions. sgRNA was extracted using phenol/chloroform and precipitated with ammonium acetate and ethanol, resuspended in 30 μl of ultrapure water (Sigma) and stored at −80 °C in 2 μl aliquots. Typically, this method produces concentrations of around 500 ng/μl; exact concentrations for each sgRNA are provided in Table 1. Working dilutions of sgRNA were made just prior to injection by diluting 3 or 5 fold. EnGen *S. pyogenes* Cas9 NLS (NEB) protein (0.3 μl) was loaded with sgRNA (1 μl for 1:3, 0.6 μl for 1:5) by incubating them together with ultrapure water for 5 minutes at 37 °C in a total volume of 3 μl. Freshly fertilised *X. laevis* eggs were de-jellied in 2% cysteine pH 7.9 and rinsed three times with 1 x MMR. Embryos were selected for injection based on the appearance of sperm entry points, and placed into a well cut into a 2% agar lined petri dish containing 6% Ficoll 400 in 1 x MMR. Cas9/sgRNA solution was loaded by backfilling into a glass capillary needle (Drummond) pulled to a fine point using a Sutter P92 needle puller and the end clipped with fine forceps. The needle was loaded onto a Drummond Nanoject II micropipette held with a MM3 micromanipulator, and embryos were injected with 9.2 nl of Cas9/sgRNA. Fifty embryos were injected at each dilution, and 50 controls were injected with only Cas9 protein. After 2 - 3 hours embryos were placed in 3% Ficoll, 0.1 x MMR. After 18 hours, they were moved to 0.1 x MMR.

**Table 1.**
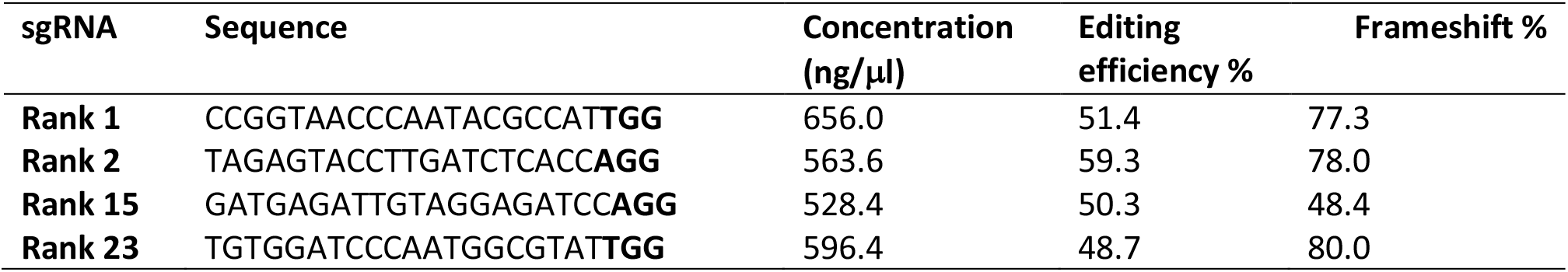
sgRNA for *Tlr4*.*S*, ranked by ChopChop v2, with PAM in bold. Stock concentrations for each sgRNA are provided, as well as predicted efficiency of editing and frameshift from InDelphi(Shen et al. 2018).

### 2.11 Genotyping and editing analysis

For cohort genotyping, eight randomly-chosen single embryos at stage 11 - 12 were collected into 0.2 μl PCR tubes and any liquid was replaced with 150 μl of 5% Chelex beads in TE (Tris/EDTA buffer, pH 8.0) with 30 μg Proteinase K. Following this, they were homogenised briefly by pipetting and incubated at 56 °C for 4 hours, then at 95 °C for 5 minutes to inactivate the enzyme. Chelex extracts were used directly for PCR and stored at 4 °C. For confirming editing in tail tips, the same process was followed except that 56 °C incubations were overnight, and vortexing was used instead of pipetting to disrupt the tissue.

PCR primers (Table 2) were as suggested by Chopchop v2 (Labun et al. 2016) for each sgRNA, amplifying approximately 250 bp around the target site. One microlitre of Chelex extracted DNA was amplified with the appropriate primers and MyTaq polymerase (Bioline) in a 20 μl volume. A T7 endonuclease I assay was used to initially confirm editing. PCR amplicons were cleaned using ExoSap-IT (Applied Biosystems) and sent for Sanger sequencing (Genetics Analysis Service, University of Otago) using the primer predicted to be furthest from the editing site. TIDE v2 (Tracking of Indels by Decomposition) (Brinkman et al. 2014) and DECODR (Deconvolution of Complex DNA repair) (Bloh et al. 2021) were used to assess the editing from the sequence trace files. An example of the editing by sgRNA rank 15 is shown in Figure S2.

**Table 2.**
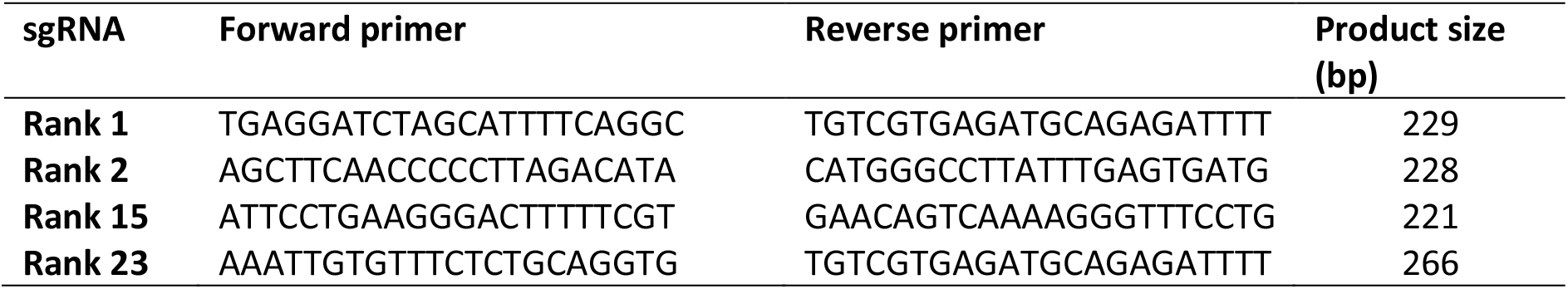
Genotyping primers for *Tlr4*.*S* Crispants.

### 2.12 Statistical analyses

Graphs were made using Graphpad Prism v9.01 or R v4.1.0 (ggplot2 (Wickham 2016)). Corresponding analyses of significant differences were performed in the same packages. Unpaired T-tests or one-way ANOVA with Tukey’s post-hoc test were used to compare the percentage of tadpoles in each dish that attempted regeneration, between untreated, antibiotic-raised and/or LPS-treated groups. Regeneration quality scores comprised of categorical data (FR, PG, PB, NR) were compared using Extended Cochran-Armitage tests or Linear x Linear association tests followed by post-hoc pairwise ordinal independence test with Benjamini-Hochberg correction for multiple testing (P.adjust). The level of CRISPR editing between regeneration categories was compared using unpaired T-test or Wilcoxon Rank Sum following Shapiro Wilk test of normality. Statistical analyses and raw data can be found in the supplemental data table.

Relative abundance plots were created in R v4.1.0 using the ggplot2 v3.3.5 (Wickham 2016) and microshades v0.0.0.9000 (Dahl et al. 2021) packages. For beta diversity analysis and visualisation, Bray-Curtis distance was calculated between samples after glomming data to genus level and normalising to relative abundance, and the vegan package (Oksanen et al. 2019) was used for permutation based ANOVA. Bacterial genera that were associated with regeneration after accounting for gentamicin use were determined by using EdgeR (Robinson et al. 2010) to fit a quasi-likelihood negative binomial generalised log-linear model with Benjamini-Hochberg false discovery correction q<0.01. Only genera seen at least 14 times in at least 20% of samples were analysed with EdgeR. The R code used for all 16S rRNA data processing and analysis is supplied at https://gitlab.com/morganx/xenopus1.

## 3. RESULTS

### 3.1 The microbiome of tadpole tail skin is consistent within, but variable between sibships, and is altered dramatically by raising tadpoles in antibiotics

Embryos were collected from three sibships and raised from the 4-cell stage in 0.1 x MMR with or without 50 μg/ml gentamicin (Figure 1A). At stage 46, 48 tadpoles from each cohort were subjected to partial tail amputation, with the tail tips collected for 16S ribosomal RNA sequencing. Regeneration was scored after 7 days. Raising embryos and tadpoles in gentamicin significantly reduced the number of tadpoles that regenerated their tails for all three sibships (Figure 1B) and also significantly decreased the quality of regeneration (Figure 1B’). Sibship accounted for 43% of microbial community variation within tails (R^2^ = 0.43, p < 0.001, PERMANOVA), while gentamicin use accounted for 14% of variation (R^2^ = 0.14, p < 0.001, PERMANOVA) (Fig 1C). Gentamicin is a broad-spectrum aminoglycoside antibiotic that targets gram-negative bacteria primarily, but not exclusively (Krause et al. 2016). Consistent with this, the bacterial population of untreated tadpole tails comprised almost entirely gram-negative taxa, while gram-positive taxa were much more abundant in gentamicin-treated tails (Figure 1D). Bacterial composition was largely consistent within sibships, but variable between sibships (Figure 1E). Sibships A and B were dominated by alphaproteobacteria, while betaproteobacteria were more abundant in sibship C. Both alpha- and betaproteobacteria were less abundant in the gentamicin-treated groups (Figure 1E).

**Figure 1:**
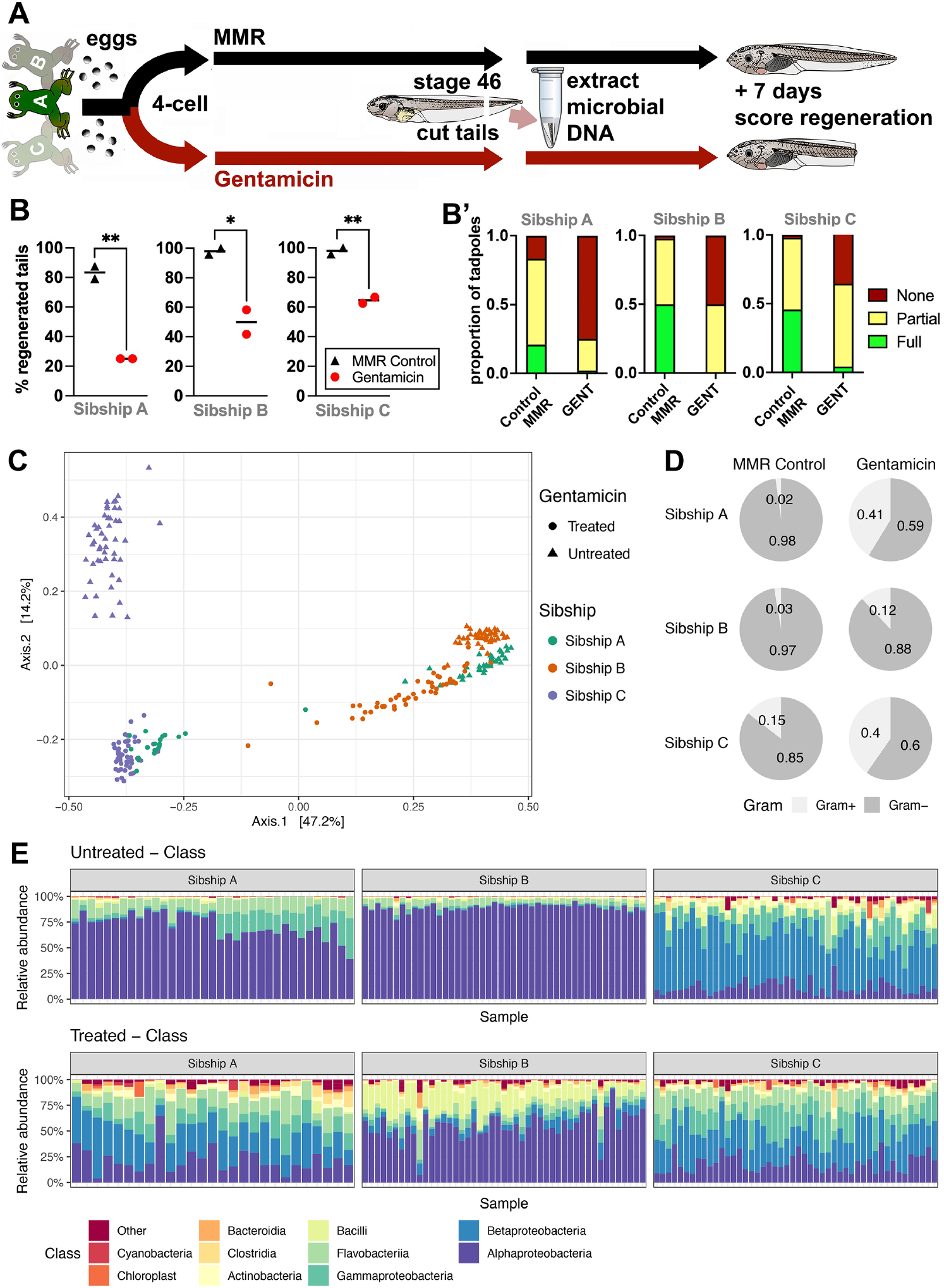
The tadpole tail skin microbiome varies between sibships and can be altered dramatically by raising tadpoles in antibiotics. **A)** Schematic of the experimental design. Three sibships of 4-cell embryos were randomly assigned to gentamicin-treated and control groups. Tail samples for microbiome analysis were obtained at stage 46 from two replicate cohorts of 24 tadpoles for each treatment and sibship. Tadpoles were scored for regeneration seven days after tail amputation. **B)** Regeneration data from three tadpole sibships. Each point represents the percentage of tadpoles regenerating any tissue at all, is the sum of full, partial good and partial bad tadpoles, and is a replicate petri dish with sample size of 24 tadpoles per dish, with the exception of controls for Sibship B where N = 22 as two died in each before they could be scored for regeneration. Unpaired t-tests, * p < 0.05, ** p < 0.01. **B’)** Stacked categorical graphs comparing regeneration phenotypes for each sibship. Linear-by-Linear association test, **** p < 0.0001. **C)** Principal coordinates analysis (PCoA) ordination plot of tadpole tail samples with > 1500 reads, calculated based on Bray-Curtis distance. **D)** Pie charts showing the percentage of gram-negative vs. gram-positive annotated reads for each sibship when raised with or without gentamicin. **E)** Relative abundance of the 10 most abundant bacterial classes in tadpole tail skin, stratified by sibship and treatment status. Raw data can be found in the supplemental data table.

We next examined how specific bacterial genera were affected by gentamicin treatment (Figure 2A), and asked if any of these were associated with successful regeneration (Figure 2B). Without treatment, each sibship was dominated by a single genus – either *Shinella* (Sibship A and B) or *Delftia* (Sibship C). Raising tadpoles in gentamicin reduced the dominance of the primary colonising genus, allowing the detection, and/or growth of, less-abundant taxa (Figure 2A). EdgeR (Robinson et al. 2010) identified six bacterial genera that were present on at least 20% of tail samples and were associated with successful regeneration (Figure 2B). These six genera varied in their relative contribution to the untreated microbial community and were generally proportionately reduced by gentamicin treatment.

**Figure 2.**
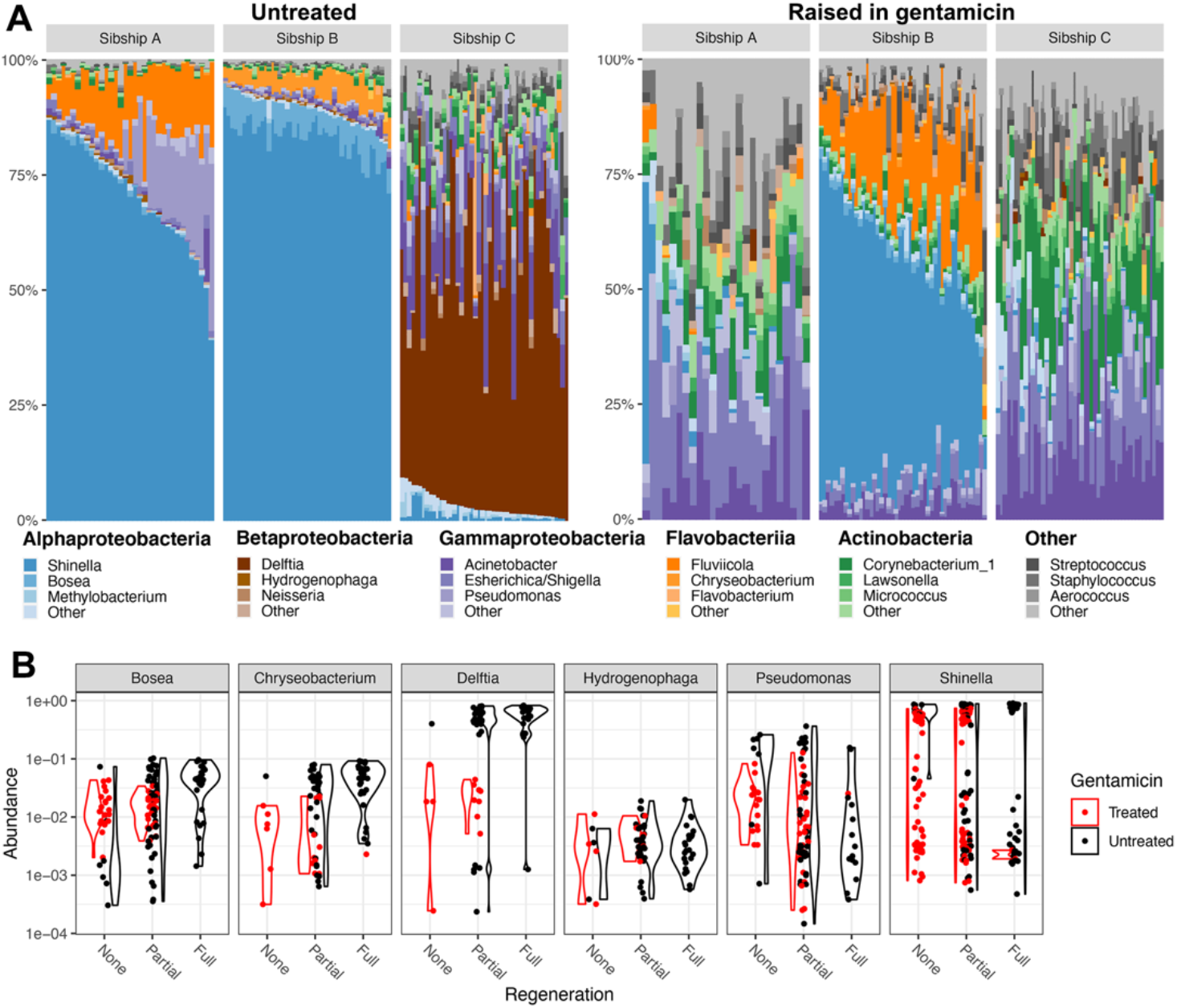
Genus-level interactions between sibship, antibiotic treatment, and regeneration. **A)** The relative abundance of genera within the five most abundant bacterial classes in treated and untreated sibships, highlighting the three most abundant genera in each. Read counts were rarefied to 1500 reads. **B)** Violin plots show log-transformed relative abundance (y-axis) of six genera positively associated with regeneration (q < 0.01, Benjamini-Hochberg false discovery correction) stratified by gentamicin status (colour). Raw data can be found in the supplementary data table.

One possible explanation for the relative increase in gram-positive taxa detected on the skin of tadpole tails when animals are raised in gentamicin is that an overall reduction of commensal bacteria allows gram-positives to bloom. To test this hypothesis, tadpoles from two further sibships were raised to stage 47 with or without gentamicin. Bacteria were recovered from the exterior surface of each tadpole and plated onto LB agar (Figure 3A). Plates inoculated from treated tadpoles generated few or no colonies, while plates from untreated tadpoles generated large numbers of colonies (Figure 3B), indicating that gentamicin was indeed effective in reducing the number of viable bacteria on tadpole skin. Control plates with no tadpole material failed to produce discernible colonies (Figure 3C). The results of these cultures suggest that overall bacterial load is reduced on the skin of these tadpoles, and the observed reduction in total number of 16S rRNA reads within normalised sequencing libraries that were generated from gentamicin treated tadpole tail samples is consistent with this (Figure S3).

**Figure 3:**
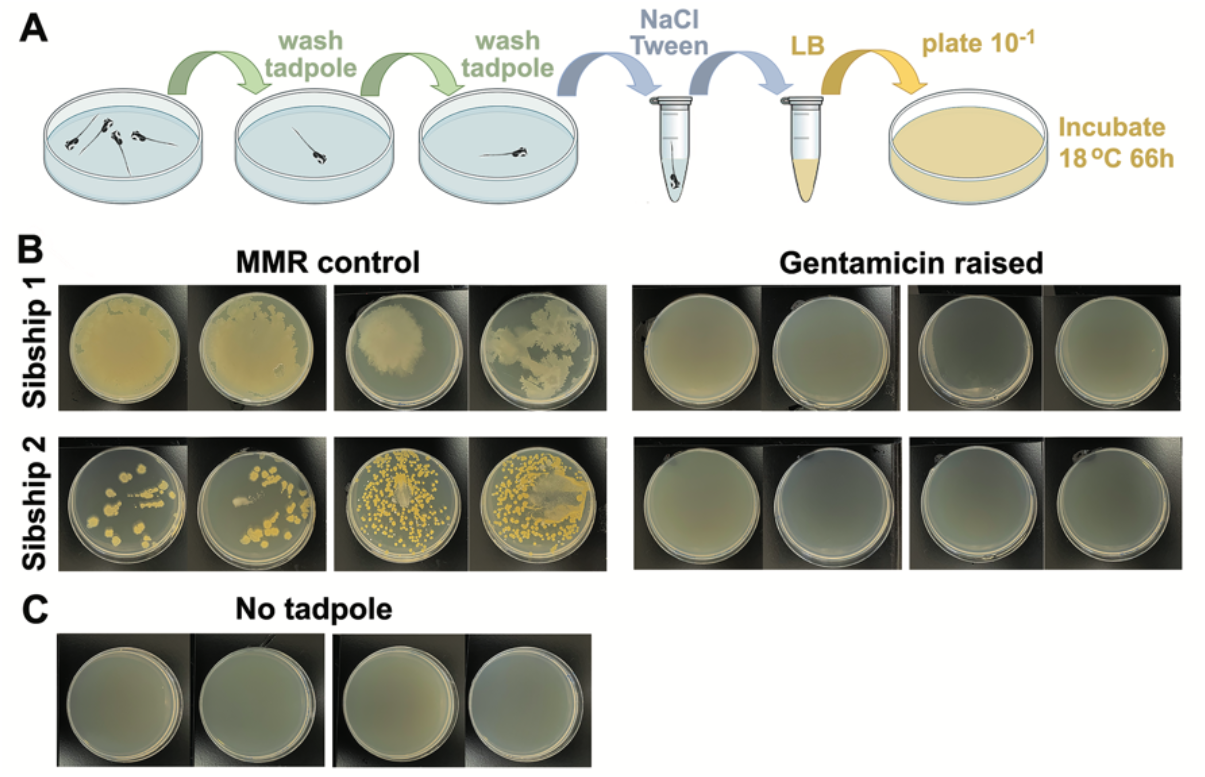
Raising tadpoles in 50 μg/ml gentamicin dramatically reduces the number of viable bacteria grown from tadpole skin. **A)** Schematic of the method used to capture bacteria from single stage 47 tadpoles. After being raised in either MMR or gentamicin solution, a selected tadpole was washed twice in MMR and vortexed for 20 seconds in 100 μl NaCl/Tween20. Fifty microlitres of the solution was then added to 1 ml Luria Broth and two replicate plates spread. **B)** Plates photographed after 66 hours at 18 °C. Two tadpoles from each sibship, raised ± gentamicin are shown. **C)** Controls prepared as above but with no tadpole, to ensure no contamination from the environment.

### 3.2 LPS from commensal *Chryseobacterium spp. XDS4* or from a *Delftia spp*. isolate can rescue regeneration in gentamicin raised tadpoles

Our previous work showed that addition of commercially-purified *E. coli* or *Pseudomonas aeruginosa* LPS to the tadpole media immediately after tail amputation rescues regeneration of antibiotic raised tadpoles to untreated levels (Bishop and Beck 2021). We hypothesised that LPS from the commensal genera that we had identified as over represented in regenerating tadpoles would also promote regeneration of refractory stage tadpoles. We adapted a method for extracting LPS from cultured bacteria, and benchmarked this against commercial preparations of *E. coli* 055:B5 LPS. Both commercial and lab-extracted *E. coli* LPS were added to gentamicin-treated tadpoles, in an attempt to rescue their regeneration ability (Figure S4A). Tadpoles from two sibships raised in gentamicin showed a significantly reduced ability to regenerate compared to untreated controls (Figure S4B-C). When added back to treated tadpoles, both forms of *E. coli* LPS were able to rescue the frequency of tadpoles undergoing tail regeneration to control levels (Figure S4B-C). The quality of the regenerates was fully rescued in one of the sibships (Figure S4 C’) but only partially in the other (Figure S4 B’).

We next attempted to isolate regeneration-associated commensal species directly from adult female *X. laevis* skin swabs. We successfully isolated two of the genera identified as regeneration biased by the differential abundance analysis (Figure 2B), a novel *Shinella* (Hudson et al. 2021) and a *Chryseobacterium spp*.. LPS was extracted from *Chryseobacterium spp. XDS4*, as *Chryseobacterium spp*. was the most overrepresented in successfully regenerating tadpoles after *Delftia spp*. (Figure 2B). The ability of *Chryseobacterium* LPS to rescue regeneration was compared to 50 μg/ml of *E. coli* 055:B5 LPS (Figure 4). In all three sibships tested, LPS from *Chryseobacterium spp. XDS4* was at least as effective as *E. coli* LPS in its ability to rescue tail regeneration following gentamicin treatment. A 50 μg/ml dose was able to restore regeneration to levels comparable with those seen in control (MMR) tadpoles, and increased doses did not result in improvement of the regeneration outcome (Figure 4B-D). For each sibship, we were able to rescue regeneration in antibiotic raised tadpoles to the level seen in control tadpoles, which varied with sibship (86%, 100%, 89% for 4B, C and D respectively). The quality of regeneration was not able to be fully rescued to control levels by LPS in one of the three sibships (Figure 4B’) but full rescues were achieved by 250 μg/ml of *Chryseobacterium* LPS (Figure 4C’) or by 50 μg/ml (Figure 4D’).

**Figure 4:**
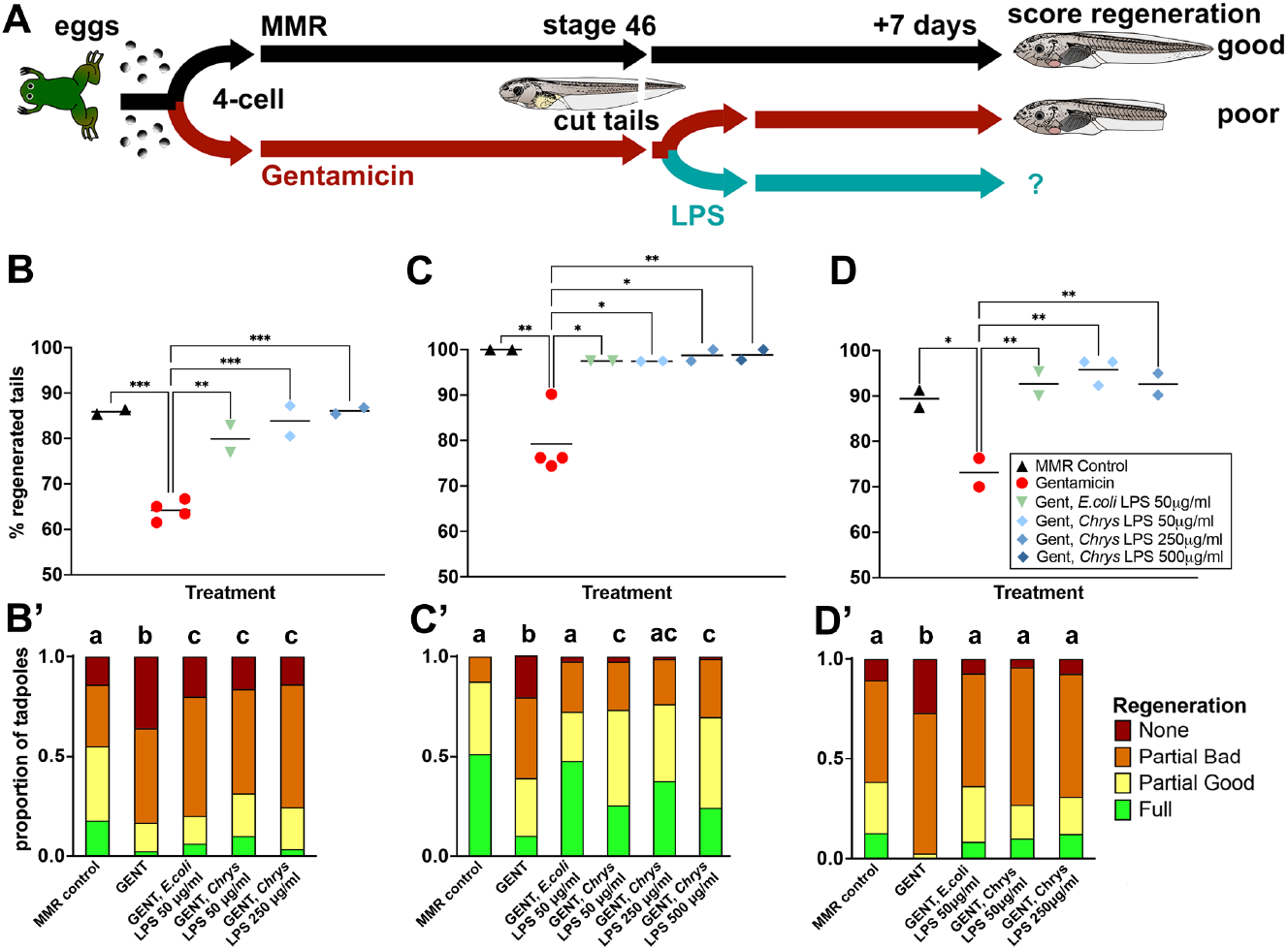
LPS from the commensal bacterium *Chryseobacterium spp. XDS4* rescues regeneration in stage 46 tadpoles raised in the antibiotic gentamicin (gent). **A)** Timeline of treatments. **B), C)** and **D)** represent data from three sibships of tadpoles. Each point represents the percentage of tadpoles regenerating any tissue at all, is the sum of full, partial good and partial bad tadpoles, and is a replicate petri dish with sample size of 38-48 (A), 35-43 (B), or 23-43 tadpoles per dish (C). 1-way ANOVA with Tukey post-hoc comparisons of all means.. * p < 0.05, ** p < 0.01, *** p < 0.001, **** p < 0.0001. **A’), B’)** and **C’)** are stacked categorical graphs of the same tadpoles, showing the proportion of each phenotype by dish. Compact letter display has been used to indicate statistical significance; each treatment is assigned a letter, with treatments within the same letter group having no statistically significant difference from each other. Extended Cochran-Armitage test followed by post-hoc pairwise ordinal independence test with Benjamini-Hochberg correction. Raw data can be found in the supplementary data table.

*Delftia* was abundant in Sibship C (Figure 2A), but we did not culture any *Delftia spp*. from frog skin. As *Delftia* was the most over-represented genus in regenerating tadpoles (Figure 2B), LPS was prepared from an isolate of *Delftia* (ICMP19763) obtained from Manaaki Whenua Landcare Research New Zealand. This LPS was found to be at least as effective as *Chryseobacterium spp. XDS4* and *E. coli* LPS at rescuing regeneration in gentamicin-raised tadpoles (Figure 5). Tadpoles from all three sibships reached control regeneration levels when raised in gentamicin and treated with *Delftia* LPS immediately post-amputation (Figure 5 A-C). Regeneration quality was also fully rescued by 50 μg/ml *Delftia* LPS in all three sibships (Figure 5A’-C’).

**Figure 5:**
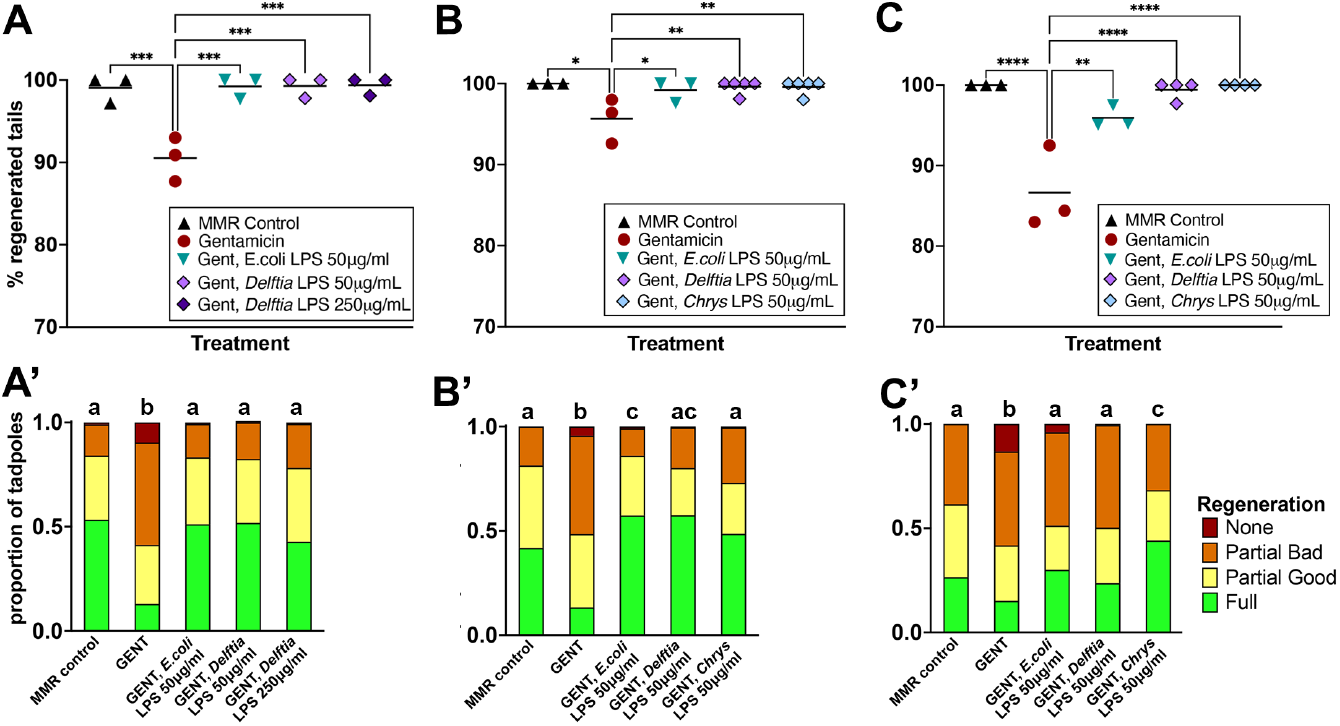
LPS from an exogenous *Delftia spp*. rescues regeneration in stage 46 tadpoles raised in the antibiotic gentamicin (gent). Timeline of treatments as for Figure 4A. **A), B)** and **C)** represent data from three sibships of tadpoles. Each point represents the percentage of tadpoles regenerating any tissue at all, is the sum of full, partial good and partial bad tadpoles, and is a replicate petri dish with sample size of 32-65 (A), 23-60 (B), or 40-65 tadpoles per dish (C). 1-way ANOVA with Tukey post-hoc comparisons of all means. * p < 0.05, ** p < 0.01, *** p < 0.001, **** p < 0.0001. **A’), B’)** and **C’)** are stacked categorical graphs of the same tadpoles, showing the proportion of each phenotype by dish. Compact letter display has been used to indicate statistical significance; each treatment is assigned a letter, with treatments within the same letter group having no statistically significant difference from each other. Extended Cochran-Armitage test followed by post-hoc pairwise ordinal independence test with Benjamini-Hochberg correction. Raw data can be found in the supplementary data table.

### 3.3 Addition of antagonistic LPS from *Rhodobacter sphaeroides* or CRISPR/Cas9 editing of TLR4 reduced regeneration quality in untreated tadpoles

We had previously suggested that TLR4 might act as the receptor for LPS (Bishop and Beck 2021), because TLR4 is the most specific PAMP (Pathogen Associated Molecular Pattern) for LPS and is known to activate the transcription factor NF-κB (Chow et al. 1999). To directly test the role of TLR4 in the regeneration pathway, penta-acetylated LPS from *R. sphaeroides*, a TLR4 antagonist (Kutuzova et al. 2001; Gaikwad and Agrawal-Rajput 2015) was added to tadpoles not exposed to gentamicin (Figure 6A). Tadpoles treated with a commercial preparation of *R. sphaeroides* LPS (Invivogen) at 50 μg/ml did not significantly reduce the percentage of tadpoles undergoing regeneration, but did significantly reduce the quality of regeneration compared with untreated controls (Figures. 6B, 6B’, Asymptotic linear-by-linear association test, *p =* 0.0428). We also prepared LPS from *R. sphaeroides* ourselves, and this was able to reduce regeneration quality to a level similar to those seen in gentamicin treated sibling tadpoles (Figure 6C & 6C’). While the standard dose of 50 μg/ml did not significantly reduce regeneration quality compared to controls, an increased dose of 250 μg/ml was able to achieve outcomes similar to those in gentamicin treated tadpoles. A further increase to 500 μg/ml did not result in any further reduction in regeneration (Figure 6C, C’).

**Figure 6:**
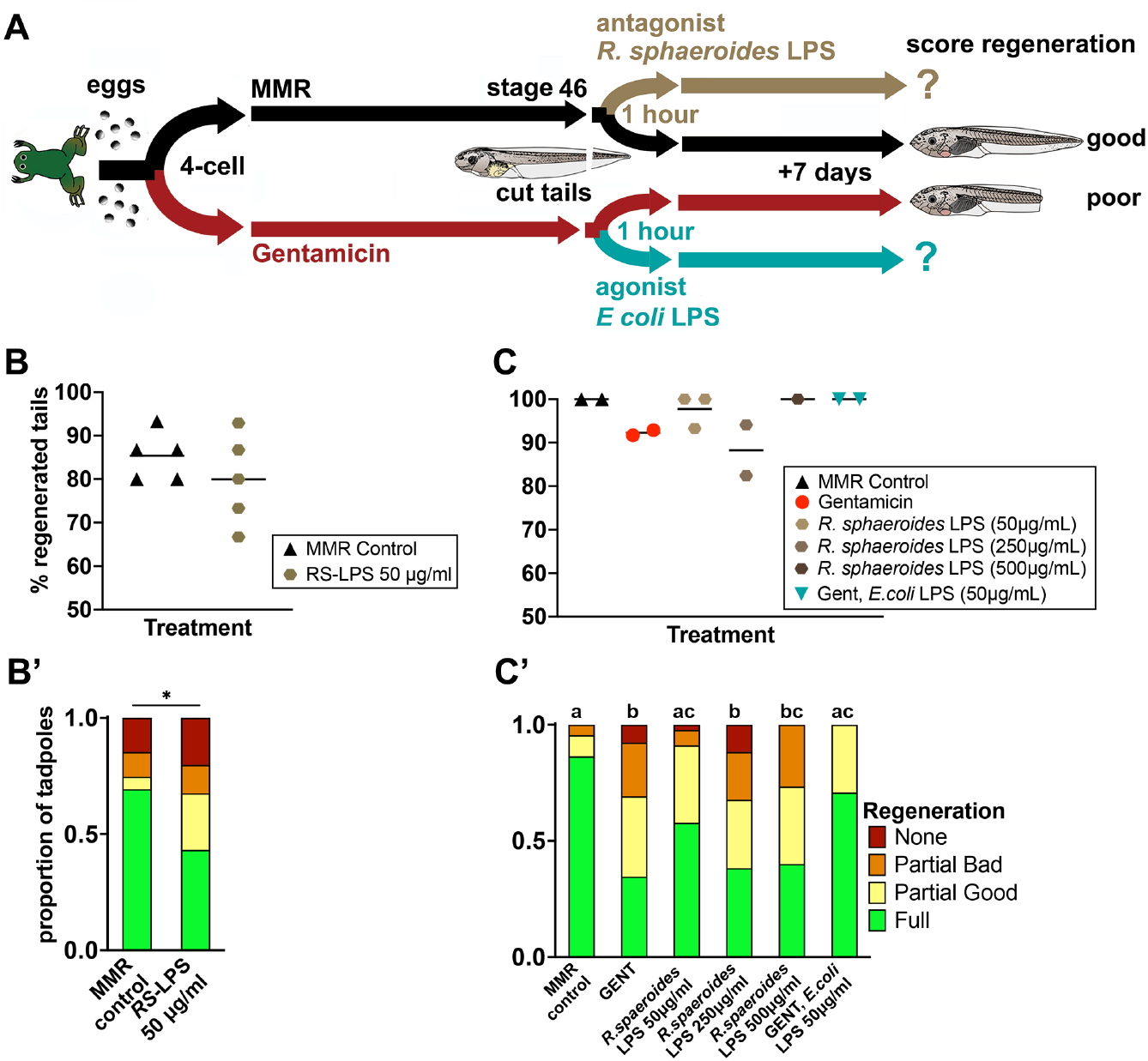
*Rhodobacter sphaeroides* LPS, a TLR4 antagonist, can significantly reduce regeneration quality, but not quantity. **A)** Timeline of treatments. Exposure of the cut tail stump to agonistic LPS should enhance regeneration in antibiotic raised tadpoles, as in Figures 2-4, and antagonistic LPS (RS-LPS) is expected to reduce regeneration in naturally raised tadpoles. **B), C)** Scatterplots where each point represents the percentage of tadpoles regenerating any tissue at all, is the sum of full, partial good and partial bad tadpoles, and is a replicate petri dish with sample size of N=15 (B) or 11-17 tadpoles per dish (C). **B)** 50 μg/ml ultrapure RS-LPS (Invivogen) treatment vs controls. Unpaired t-test showed no significant difference between groups. **C)** Post-amputation treatment with three concentrations of antagonistic extracted RS-LPS were compared to control tadpoles and gentamicin treated tadpoles with or without *E. coli* LPS rescue. 1-way ANOVA with Tukey post-hoc comparisons of all means showed no significant differences between groups. **B’)** and **C’)** are stacked categorical graphs of the same tadpoles, showing the proportion of each phenotype by dish. Compact letter display has been used to indicate statistical significance; each treatment is assigned a letter, with treatments within the same letter group having no statistically significant difference from each other. Extended Cochran-Armitage test followed by post-hoc pairwise ordinal independence test with Benjamini-Hochberg correction. * p < 0.05. Raw data can be found in the supplementary data table.

As a second approach, we used CRISPR/Cas9 to disrupt the *Tlr4*.*S* gene. *Xenopus laevis* is allotetraploid(Session et al. 2016), but there is only a single copy of TLR4. We predicted that editing would lead to gene function disruption and a subsequent reduction of regeneration in crispants. Four sgRNAs were designed and trialled to determine their efficiency in editing *Tlr4*.*S* (Figure 7A, Table 1 and 2). Of these, sgRNA rank 15 at a 1:3 concentration, predicted to cause a frameshift resulting in a premature stop codon (Figure 7A), was the only sgRNA to achieve a high level of editing in embryos (74%, Figure 7B, Figure S2). The effect of *Tlr4*.*S* editing on frequency of tadpole tail regeneration was not significant, but the quality of regeneration in crispants was significantly lower than for controls (Extended Cochran-Armitage test, *p =* 0.000000132). In the same sibship, *R. sphaeroides* LPS also reduced regeneration quality (p < 0.0001). Gentamicin-raised tadpoles had significantly lower levels of regeneration than *R. sphaeroides* treated or crispant tadpoles (Figure 7B). Sequence analysis of embryos using the three other sgRNA, or with Cas9 alone, showed no significant editing, and regeneration quality was indistinguishable from controls (Figure 7B). Taken together, we suggest that partial inhibition of TLR4 signalling, by either excess antagonist LPS or partial gene editing with CRISPR/Cas9, does reduce the quality of tadpole regeneration.

**Figure 7:**
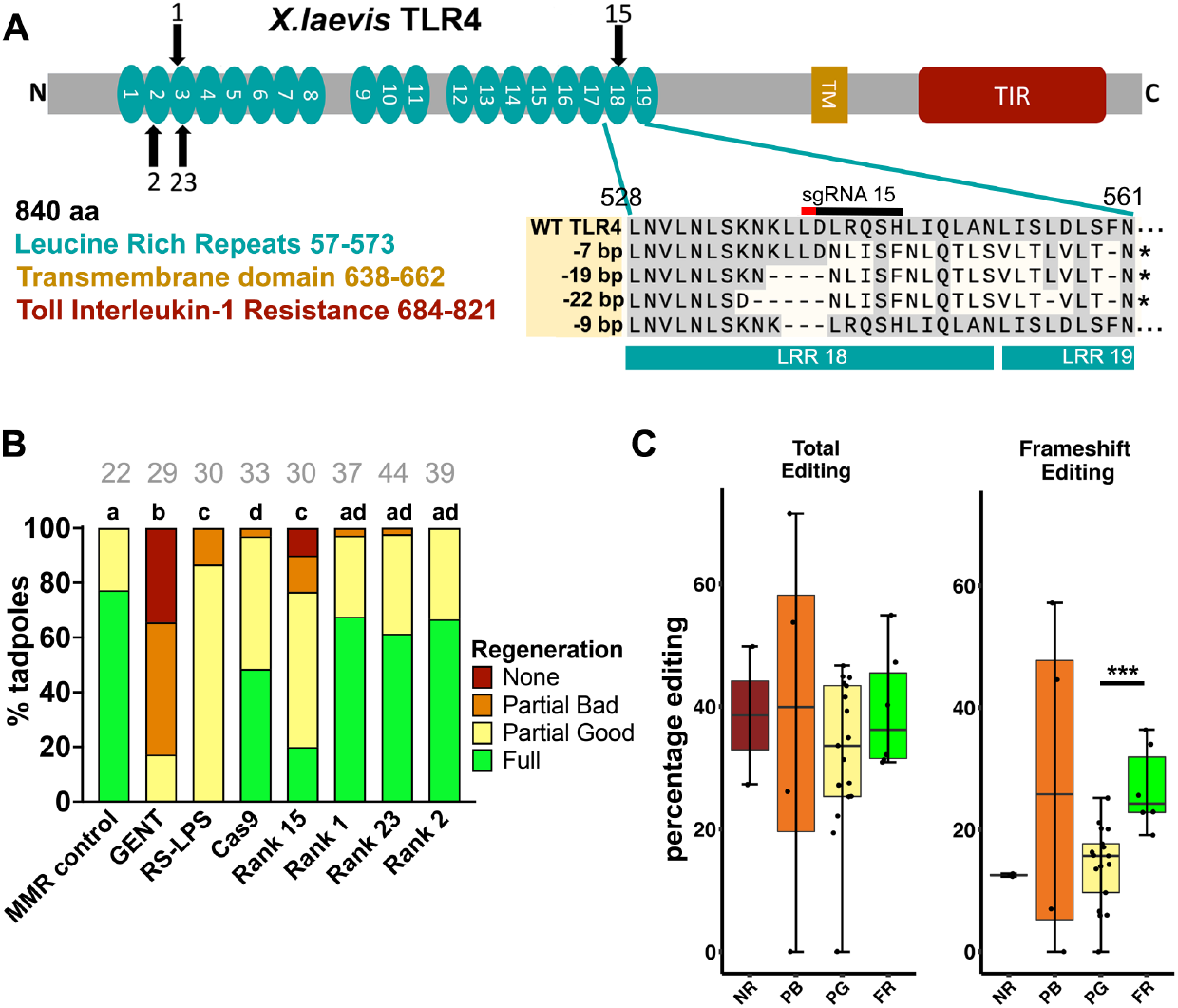
TLR4 editing with CRISPR/Cas9 correlates with reduced regeneration score at sibship but not individual tadpole level. **A)** Schematic of *X. laevis* TLR4 protein, showing 19 predicted extracellular LRR domains, an internal Toll-interleukin-1 inhibition domain (TIR) predicted by NCBI CDD and a single transmembrane domain (predicted by TMHMM server v2.0)(Sonnhammer et al. 1998). Black arrows show targets of sgRNA, numbers associated with arrows indicate the specific sgRNA. The most common deletions generated by sgRNA rank 15 result in a +1 frameshift which leads to a stop codon that truncates the protein mid 19^th^ LRR domain. A less commonly occurring −9 bp deletion results in the loss of three amino acids from LRR 18. **B)** Stacked categorical graphs of tadpole regeneration, showing the proportion of each phenotype by treatment. MMR controls are unmanipulated embryos, gentamicin is embryos raised in 50 μg/ml gentamicin from 4 cell stage to 1 day post amputation. No other embryos in this set were raised in antibiotics. RS-LPS is *R. sphaeroides* LPS, a natural TLR4 antagonist. Four different sgRNA were used, average editing for eight randomly chosen stage 10 embryos was 74% for sgRNA rank 15 and < 10% for the other sgRNA. Tadpoles at stage 46 had the posterior third of the tail removed using a scalpel blade and were scored for regeneration quality a week later. Compact letter display has been used to indicate statistical significance; each treatment is assigned a letter, with treatments within the same letter group having no statistically significant difference from each other. Grey numbers above bars indicate sample size (N). Extended Cochran-Armitage test followed by post-hoc pairwise ordinal independence test with Benjamini-Hochberg correction. **C)** Boxplot of total editing percentage (C) and frameshift editing percentage (C’) in tadpole tail clips from each regeneration category. Shapiro-Wilk test of normality followed by unpaired T-test (Editing) and Wilcoxon Rank Sum (Frameshift). NR = No Regeneration, PB = Partial Regeneration (Bad), PG = Partial Regeneration (Good), and FR = Full Regeneration. *** p < 0.001. Raw data can be found in the supplementary data table. * p < 0.05, ** p<0.01, ***p<0.001. Raw data can be found in the supplementary data table.

Individual tadpole tail clips from the sgRNA rank 15 group were also checked for editing, which ranged from 19 to 54% across 27 individuals. Mean editing in the cohort was 34%, with frameshift editing at just 17.8%. To see if individual editing levels influenced the regenerative outcome of tadpoles, we compared editing levels in tadpoles grouped by the four regeneration categories. No significant difference in either total editing or frameshift editing percentage was evident between the groups (Figure 7C) except when frameshift editing in the FR and PG groups were compared. Overall, this indicates that a particular tadpole was not less likely to regenerate if its *Tlr4*.*S* editing level was higher.

## 4. DISCUSSION

Amphibian tadpoles, like all metazoa, support populations of microorganisms that interact with their hosts through various mechanisms. Here, we show that the tadpole skin microbiome is highly variable and can be manipulated by raising embryos in the antibiotic gentamicin. Six gram-negative genera, including *Delftia* and *Chryseobacterium*, were over-represented in tadpoles that successfully regenerated their tails. Regeneration could be rescued in antibiotic-raised tadpoles by adding LPS from commensal *Chryseobacterium spp. XDS4, Delftia Wen et al 1999*, or *E. coli*. Conversely, regeneration was impaired in tadpoles exposed to an antagonistic LPS isolated from *R. sphaeroides*. Disrupting *Tlr4*.*S* using CRISPR/Cas9 also reduced regeneration quality, but not quantity, at the level of the cohort. However, we found that the editing level of individual tadpoles was not a good predictor of regenerative outcome.

### 4.1 The *X. laevis* pre-feeding tadpole skin microbial community varies with sibship, lacks diversity, and can be manipulated with antibiotics

Gram-negative bacteria, in particular Proteobacteria, were dominant over gram-positive phyla in the tadpoles’ unmodified microbiome (Figure 1). However, the dominant bacterial clades varied between sibships; the alphaproteobacteria class was predominant in two sibships (A and B), while betaproteobacteria dominated the third (C). Sibship B had highest detected levels of alphaproteobacteria, and retained these at higher levels than in sibships C or A, when raised in gentamicin. This variation between tadpole cohorts may be partly due to host genetics, but is probably also attributable to environmental factors. While *Xenopus* microbiome work is in its infancy, Piccini et al. (2021)(Piccinni et al. 2021) found that although the adult *X. laevis* skin microbiome is subject to strong selective pressures from the host, tadpole microbiomes were more variable and influenced by environmental conditions. Interestingly, the microbiomes of the older, premetamorphic tadpoles in the Piccini study(Piccinni et al. 2021) were also dominated by proteobacteria, although were not dominated by single genera as ours were. However, this is almost certainly affected by differences in stage/age and sample collection methods. Piccini et al. swabbed month old tadpoles (expected stage 54-55, length 60-80 mm) that had been fed algae and were housed in aquaria(Piccinni et al. 2021), whereas those in our study were maintained in petri dishes in MMR at a constant 18 °C, were approximately 6-7 days old (stage 46, 9-12 mm length) at sampling, and had never been fed. Further, Piccini’s tadpoles and frogs were routinely raised for the first week in penicillin and streptomycin, and could therefore have acquired their microbiome from tank water and food(Piccinni et al. 2021). The comparison between our study and that of Piccini et al. demonstrates that *Xenopus* microbiomes undoubtedly vary from one laboratory to another based on husbandry and other environmental variables. Until now, no *Xenopus* regeneration studies have accounted for variations in microbiomes. In order to understand how commensals influence tail regeneration, it will be important in the future to determine both the source of the *Xenopus* microbiome and how it evolves at various life stages as a critical step in determining the relative contribution of microbes and genetics.

### 4.2 Gram-negative LPS concentrations and/or specific genera may determine the regenerative response

As expected, raising tadpoles in a gentamicin solution resulted in altered microbiome composition, increased the proportion of gram-positive bacteria, and decreased regeneration success compared with untreated tadpoles. While we cannot entirely eliminate the possibility that gentamicin treatment introduces an anti-regenerative effect separate to the reduction of pro-regenerative gram-negative bacteria, successful rescues through the addition of LPS to the antibiotic media suggests that the antibiotics cause no significant disruption to other facets of the regeneration pathway.

Based on our results and those our previous studies omitting antibiotics (Taylor and Beck 2012), (Bishop and Beck 2021), baseline regeneration rates among untreated refractory stage 46 tadpoles appear to be variable between sibships, ranging from 55 to 100%. It is unclear whether this is due to genetic factors, environmental factors, the presence of anti-regenerative microbial taxa in the microbiome, or an interplay of all three. Abundance of *Rhodobacter spp*., demonstrated here to have an anti-regenerative effect, was low (just 333 reads in total across all samples). It remains to be investigated whether other taxa observed could have a similar effect. It is difficult to compare our baseline regeneration rates with those from others’ studies, as parentage and antibiotic exposure of tadpoles is not always declared.

Six bacterial genera were more abundant on the skin of successful regenerators: *Pseudomonas, Bosea, Shinella, Chryseobacterium, Delftia* and *Hydrogenophaga*. Previously, we showed that a commercial preparation of *P. aeruginosea* LPS restores tail regeneration ability in antibiotic-raised stage 46 tadpoles (Bishop and Beck 2021). Here, we showed that LPS isolated from *Chryseobacterium spp. XDS4* and *Delftia spp*. were also able to rescue the regeneration process in gentamicin raised tadpoles. While we cannot rule out that innate features of the LPS from these particular taxa specially facilitates regeneration pathways, it seems unlikely, as *E. coli* LPS is also equally effective (Bishop and Beck 2021) (Figure S4). It is possible the total LPS load from any gram-negative commensal (with the exception of divergent, antagonistic LPS) is sufficient to determine regenerative success in the refractory period.

In this study, *Chryseobacterium spp. XDS4*, from which LPS was obtained, was cultured from adult frogs. However, it is unclear from 16S rRNA data whether this isolate is identical to the *Chryseobacterium* detected on tadpoles. None of the six genera of note identified here, with the exception of *Pseudomonas*, was found among the top 50 genera detected on tadpoles or adults in the recent Piccini et al. study (Piccinni et al. 2021), which is to date the only other such report of skin microbiota in *Xenopus* tadpoles. Although the data suggested that tadpole skin microbiomes are shaped environmentally, a lack of parental contribution was not directly determined. Here, we show that the very early tadpole microbiome is dominated by proteobacteria, and that different sibships can have different genera dominating their microbiome.

The mean number of sequencing reads collected for gentamicin-treated samples was lower than for untreated samples in all sibships, but this was most pronounced in Sibship A. We sequenced DNA from 50 tadpoles per sibship / treatment, and DNA quantities were standardized by the sequencing facility both during sequencing library preparation and final pooling prior to sequencing. However, library preparation was unsuccessful for approximately one third of gentamicin-treated Sibship A samples. The DNA in this study was extracted from whole tail samples, and is thus a mixture of tadpole and microbial DNA in proportions that may vary between samples. The lower numbers of reads generated from the DNA of gentamicin-treated samples is consistent with a reduction of total bacterial numbers in gentamicin-treated tadpoles, with a consequent decrease in dominant gram-negative bacteria and their LPS. Further support comes from the much higher numbers of colonies obtained from tadpole skin extracts when gentamicin was not used, although this used two different sibships. A quantitative assessment of LPS could be done in future to test the correlation more directly.

### 4.3 Commensal microbiota may have a critical role in regeneration and scar free wound healing

While the role of individual taxa is a developing area of research, recently, evidence is emerging to support a critical role for the microbiome in regeneration and wound healing in other model organisms. In *Schmidtea mediterranea*, free living flatworms with remarkable regeneration abilities, a pathogenic microbiome has been shown to shown to derail regeneration (Arnold et al. 2016). *Aquitalea sp*. FJL05, a gram-negative commensal bacterium of another planarian, *Dugesia japonica*, can dramatically affect the pattern of regeneration, resulting in worms with two heads (Williams et al. 2020). In this case however, indole, a small molecule produced by *Aquitalia*, rather than LPS, was the cause of the effect. Two recent studies highlight the potential role of microbiota in mouse skin and ear regeneration. Wang et al (2021) reported that germ-free mice showed reduced levels of wound-induced hair follicle neogenesis and stem cell markers. The inflammatory cytokine IL-1β and keratinocyte-dependent IL-1R-MyD88 signalling was found to be essential for regeneration (Wang et al. 2021). In healer MRL mice, Velasco et al (2021), showed that healing of ear punch wounds is linked with the gut microbiome. Excitingly, this healing ability could be transferred to non-healer mice by faecal transplant (Velasco et al. 2021).

### 4.4 TLR4 signalling may contribute to the regenerative response in tadpole tails

TLR4 signalling is not as well characterised in amphibia as it is in mammals. Recent work in urodele amphibia (axolotl) showed that inflammatory responses to PAMP ligands, such as LPS, through TLRs, are conserved. However, responses to Damage Associated Molecular Patterns (DAMPs) were found to have fundamental differences from those seen in mammals (Debuque et al. 2021). We note that orthologs of CD14 and MD2, which in mammals aid in the presentation of LPS to TLR4, appear to be absent from the *Xenopus* genomes. A third regulator of this interaction, lipopolysaccharide binding protein (LBP), is present, and would be worth targeting in future.

Our results partially support the involvement of LPS-TLR4 in regenerative pathways suggested in Bishop and Beck (2021). Addition of LPS from *R. sphaeroides*, a known TLR4 antagonist) to antibiotic raised tadpoles lead to reduced regeneration quality, with a similar effect seen in *X. laevis Tlr4*.*S* crispants. However, the inhibition of regeneration in these experiments was not absolute. *Rhodobacter sphaeroides* LPS, while achieving significant quality reduction, was not able to completely suppress regeneration, possibly due to competition for binding sites from remaining TLR4 agonist microbes. As discussed earlier, very few *Rhodobacter* sequences were detected, suggesting that *R. sphaeroides* is unlikely to be physiologically relevant in tadpole regeneration. In the CRISPR/Cas9 experiments, 100% editing was not achieved for any tadpole, despite trialling multiple sgRNAs and sgRNA concentrations to maximise efficacy. Mosaicism is an inherent problem with CRISPR/Cas9 editing and results in unedited cells within an embryo, potentially leaving a proportion of TLR4 signalling pathways intact. This would at least partially account for the persisting (albeit qualitatively poorer) regeneration capability in tadpole cohorts. Additionally, the multiple potential edits produced by any given sgRNA are unlikely to be equal in their effect on gene function (e.g. frameshifts vs. in frame InDels). These factors taken together may go some way to explaining the lack of correlation between editing percentage and rehabilitation outcome in individual tadpoles, despite a significant correlation for the cohort taken as a whole. While direct injection of the sgRNA-Cas9 protein complex minimises mosaicism over delivering DNA plasmids encoding sgRNA/Cas9(Burger et al. 2016), strategies such as simultaneous use of multiple sgRNAs (Zuo et al. 2017) and crossing of F_0_ crispants to generate complete knockouts in F_1_ (Feehan et al. 2017) could be used in future to knock out *Tlr4*.*S* completely. Further to the above, it has been demonstrated that gene knockout can lead to upregulation of related genes in compensation (El-Brolosy et al. 2019).

Theoretically, this would dampen the effect of TLR4 knockdown and allow some level of regeneration to proceed. A recent CRISPR/Cas9 knockdown of TGFβ1, one of the earliest players known to be required for tail regeneration(Ho and Whitman 2008) using three sgRNA also demonstrated a reduced quality, delayed tail regeneration response in *X. tropicalis* (Nakamura et al. 2021). TLRs have broad specificity to detect PAMPs and each receptor has its own ligand preference(Akira and Takeda 2004). While TLR4 plays a central role in mediating responses to LPS, it is possible that LPS also stimulates other receptors. TLR2 may also be responsive to LPS (reviewed in (de Oliviera Nascimento et al. 2012), and so it may be necessary to target TLR2 and TLR4 together to prevent LPS signalling. TLR4 can also be activated by DAMPs (Piccinini and Midwood 2010), such as heat shock protein HSP60 (associated with regeneration in fish (Makino et al. 2005) and frogs (Pearl et al. 2008) as well as extracellular matrix components like heparan sulphate (associated with amphibian regeneration (Phan et al. 2015; Wang and Beck 2015)) and tenascin C. A future approach could be to edit the gene for lipopolysaccharide binding protein (*Lbp*.*L*), which may mediate TLR4 receptor-LPS ligand binding. The cytoplasmic adaptor MyD88 has been implicated in axis formation in early development of *Xenopus* (Prothmann et al. 2000), and so is not a usable target.

### 4.5 Conclusions

Our results demonstrate that LPS from gram-negative bacteria enhances regenerative outcomes in *X. laevis* tadpoles, and that the signalling pathway mediating this response involves TLR4, at least in part. We suggest that future studies should examine the concurrent roles of other candidate receptors using gene knockdown, and also survey the individual effects of LPS from a broad range of bacterial taxa. Ultimately, this line of study has the potential to improve medicinal and veterinary outcomes in wound healing and regeneration.

## ACKNOWLEDGEMENTS

We are grateful for funding from Royal Society of New Zealand Marsden Fund 19-UOO-245 to support this work. CG was partly funded by a University of Otago Master’s research scholarship.

## AUTHORSHIP

Conceptualization: CB, XM. Data curation: DH. Formal Analysis: PC, CB, XM. Funding acquisition: CB, XM. Investigation: TD, CG, DH, CB. Methodology: CB, JW, XM, CG. Project administration: XM, CB. Supervision: XM, CB. Visualization: CB, XM, PC, DH Writing – original draft: PC, CB, XM Writing – review & editing: PC, CB, XM

## SUPPLEMENTARY FIGURES

**Figure S1:**
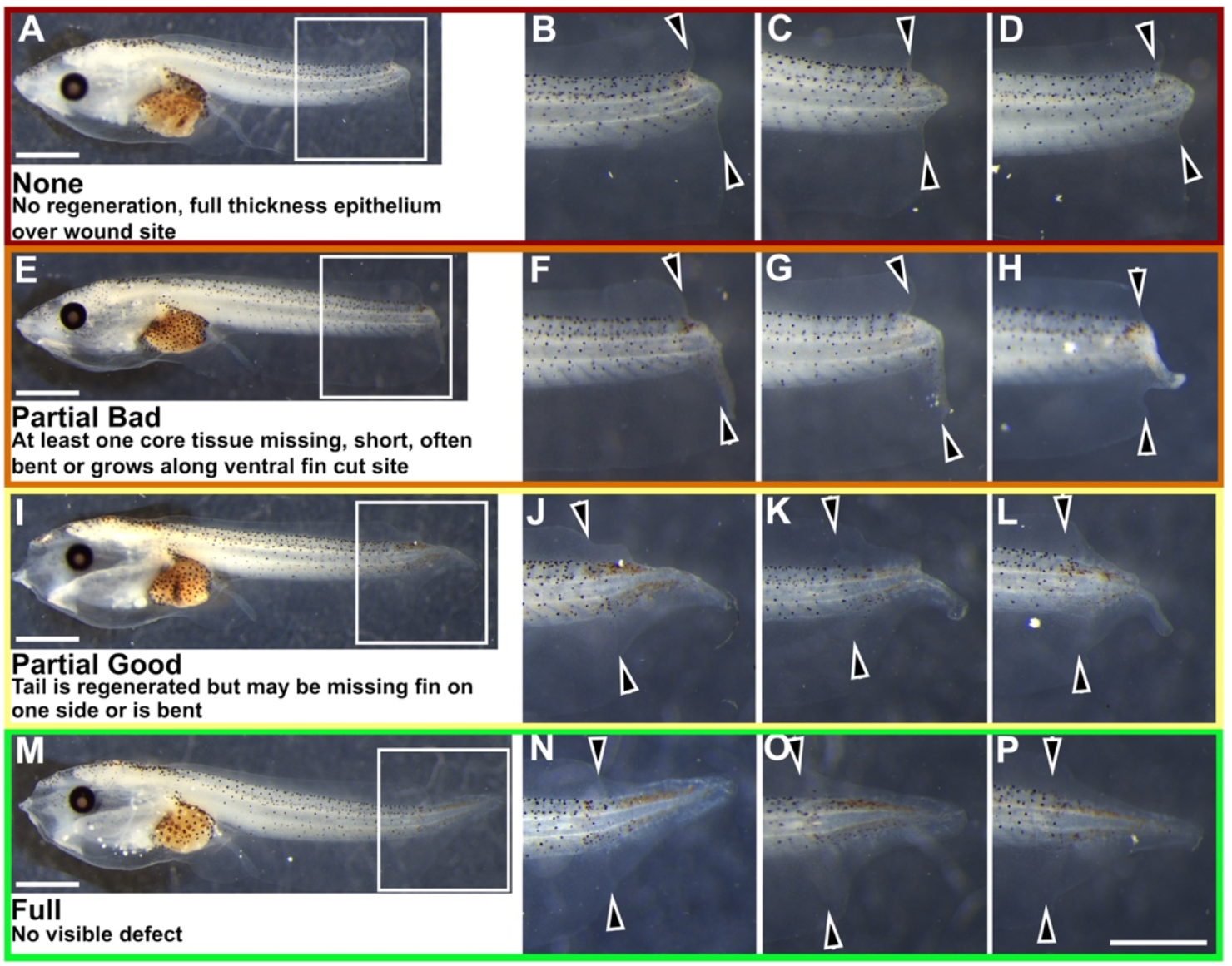
Representative examples of tail regeneration categories. Three examples for each category are shown. Tadpoles are all from Sibship C in Figure 1. Boxes in A, E, I, M relate to panels B, F, J and N respectively, and scale bars are 1 mm. Scale bar in P is 1 mm and applies to all tail panels. Arrowheads indicate the level of the tail amputation at stage 46. A-D No regeneration, wound heals, E-H Partial bad regeneration, I-L Partial good regeneration, M-P Full regeneration. For Figure 1, Partial bad and partial good were binned together as “partial”. Where no regeneration occurs (A-D), a full thickness wound epithelium forms, the bulge of axial tissue results from notochord cells, which have large vacoules for structural support (Beck et al. 2003).

**Figure S2:**
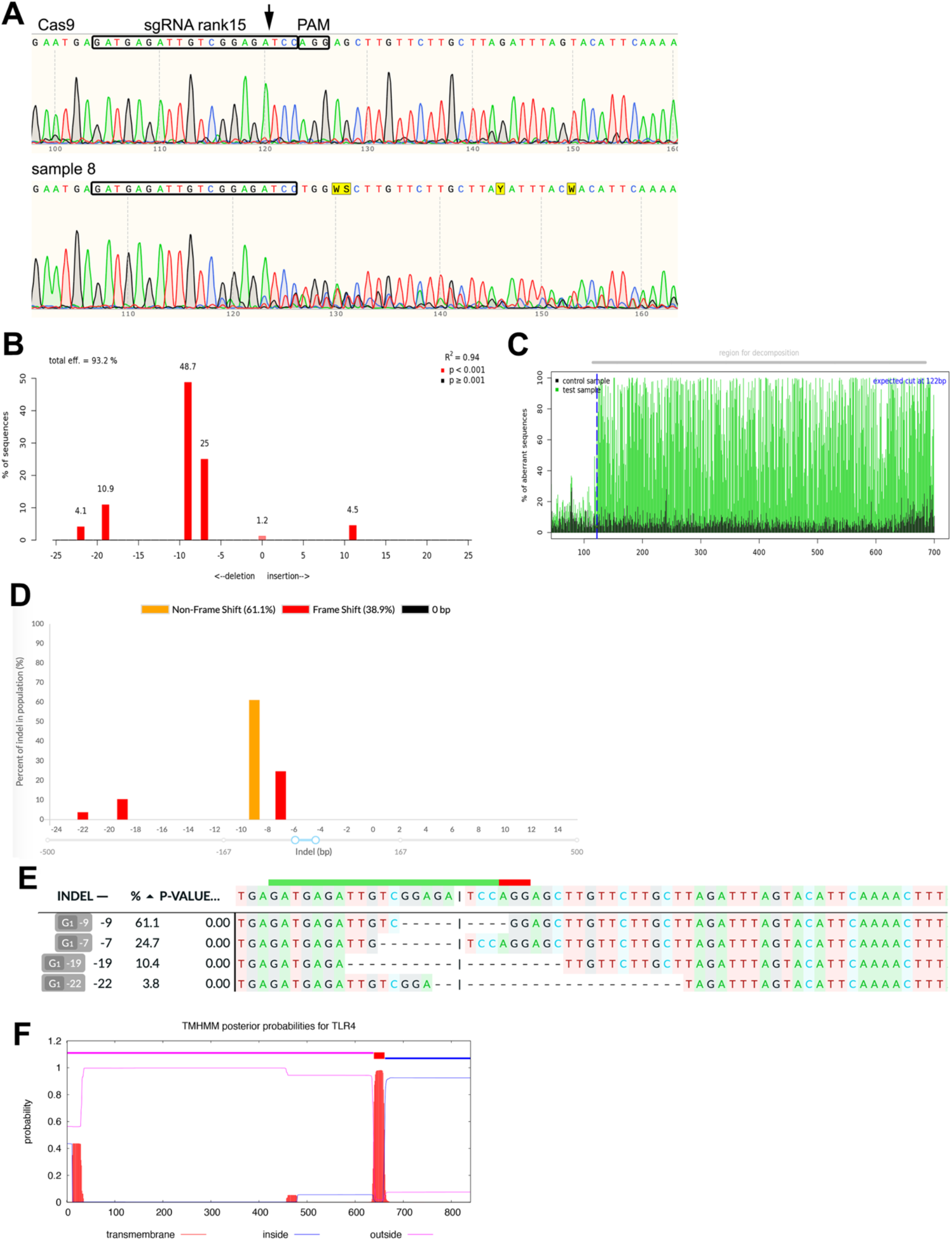
Supporting data for *Tlr4*.*S* gene editing in *Xenopus laevis*, examples of sequence analysis and editing consequences. **A)** Sanger sequencing traces from a control (cas9 injected) and edited (cas9 and sgRNA rank15) embryo from the experiment shown in Figure 7 of the main manuscript. The traces from **(A)** analysed using TIDE (Brinkman et al. 2014) **(B, C)** and Decodr v3 (Bloh et al. 2021). **(D, E). F)** The likely location of the *X*.*laevis* TLR4 transmembrane domain as calculated using TMHMM v2.0 (Sonnhammer et al. 1998).

**Figure S3.**
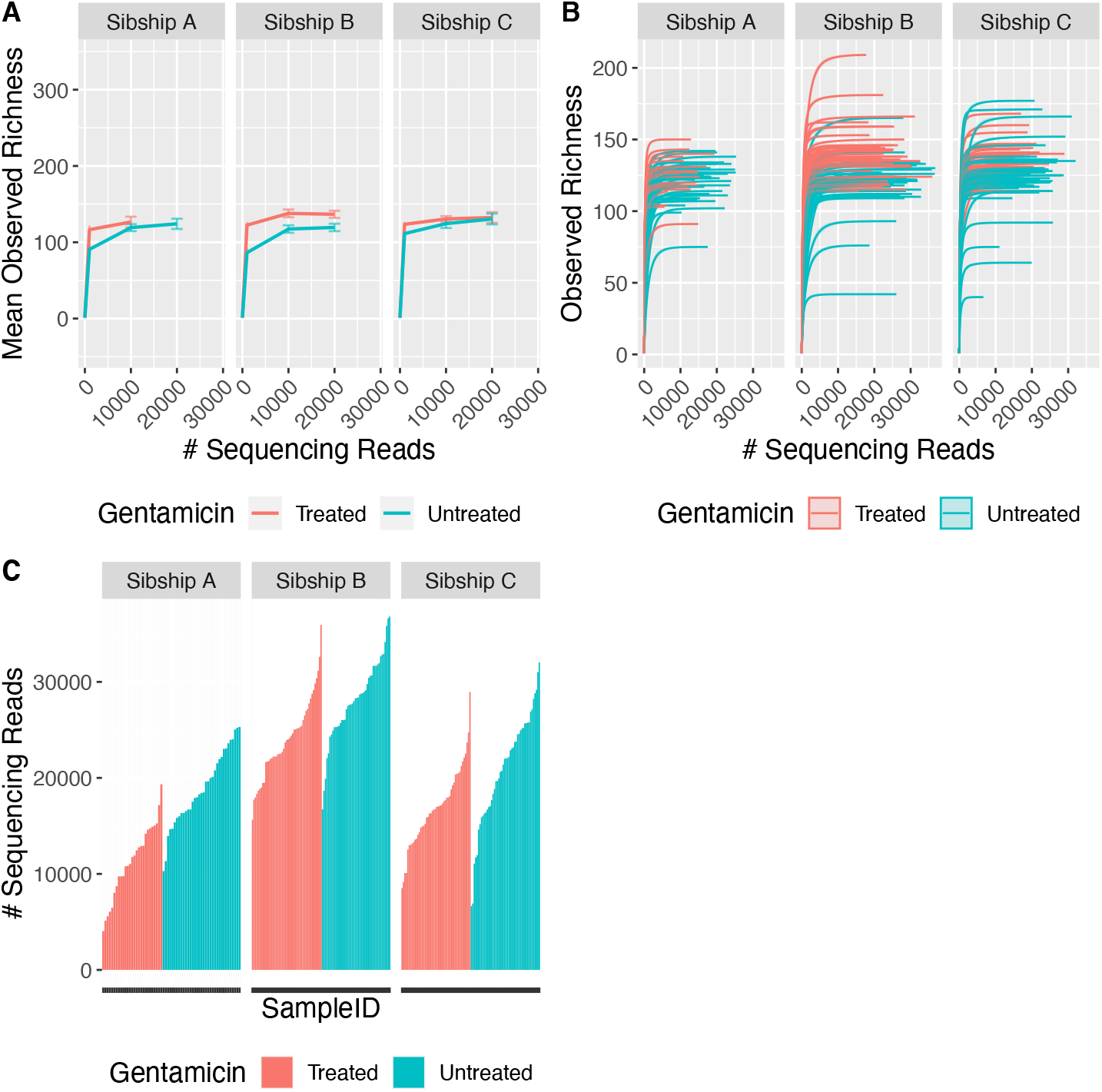
**A)** Rarefaction curve, showing mean observed alpha diversity relative to sequencing depth. Error bars correspond to 95% confidence interval for standard error mean. Samples are stratified by sibship and treatment. B) Rarefaction curve, showing observed alpha diversity relative to sequencing depth for all samples. Samples are stratified by sibship and coloured by treatment. C) Histogram summarizing total sequencing reads for each sample, grouped by sibship and coloured by antibiotic treatment status. Number of tail samples sequenced for each group was as follows. Sibship A: untreated 35, treated 27. Sibship B: untreated 44, treated 48. Sibship C untreated 47, treated 48.

**Figure S4:**
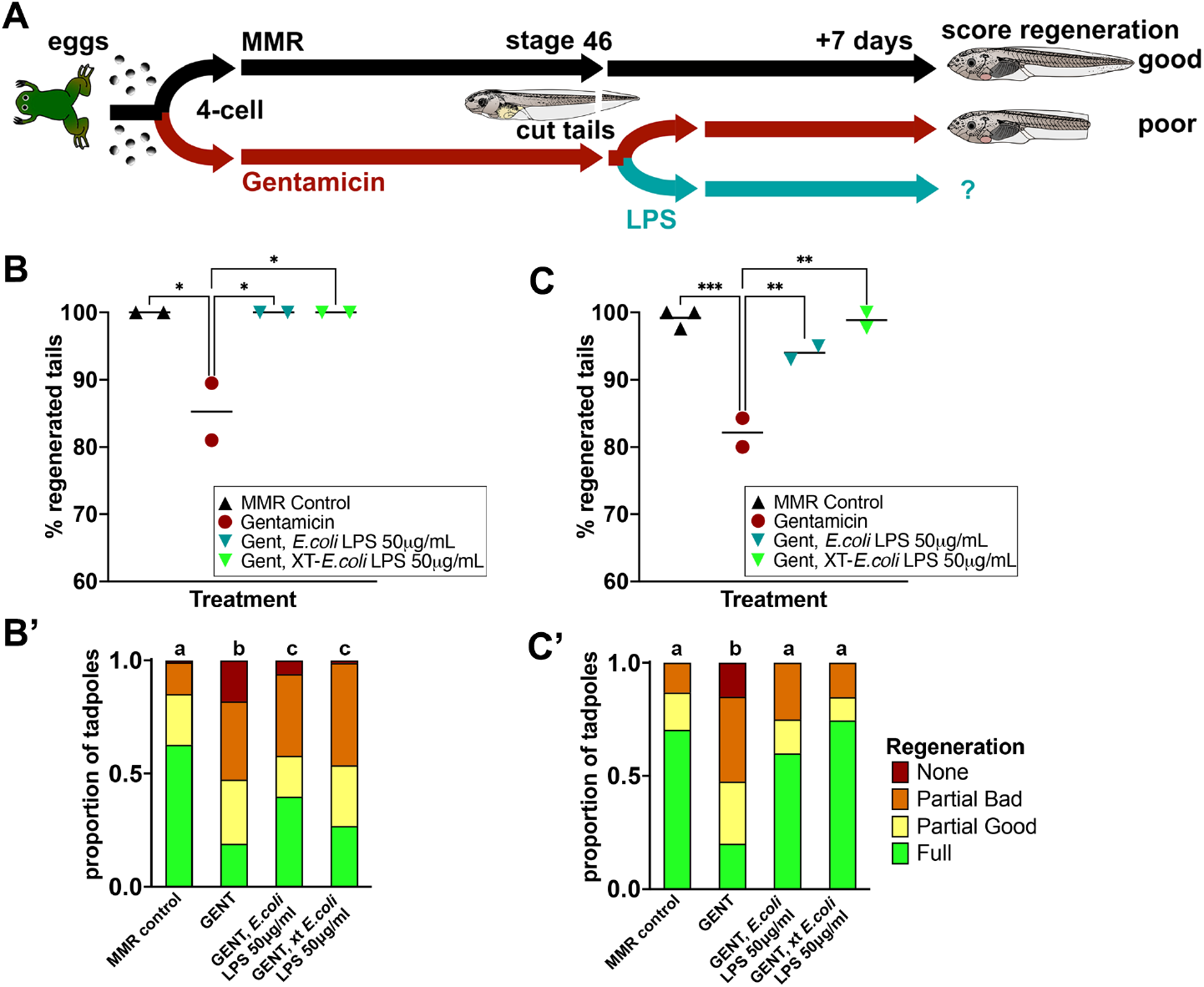
Commercial *E. coli* LPS 055:B5 (Sigma) and extracted (xt) *E. coli* LPS rescue regeneration in stage 46 tadpoles raised in the antibiotic gentamicin (gent). **A)** Timeline of treatments. **B)** and **C**) represent data from two sibships of tadpoles. Each point represents the percentage of tadpoles regenerating any tissue at all, and is the sum of full, partial good and partial bad categories in a replicate petri dish with sample size of 39-65 **(B)** and 19-46 **(C)** per dish. **B**’) and **C’)** are stacked bar graphs of categorical data pooled for each treatment, showing relative abundance of each phenotype. Compact letter display has been used to indicate statistical significance; each treatment is assigned a letter, with treatments within the same letter group having no statistically significant difference from each other. Extended Cochran-Armitage test followed by post-hoc pairwise ordinal independence test with Benjamini-Hochberg correction. * p < 0.05, ** p<0.01, ***p<0.001. Raw data can be found in the supplementary data table.

